## Supplemental Figures for "High throughput machine learning pipeline to characterize larval zebrafish motor behavior"

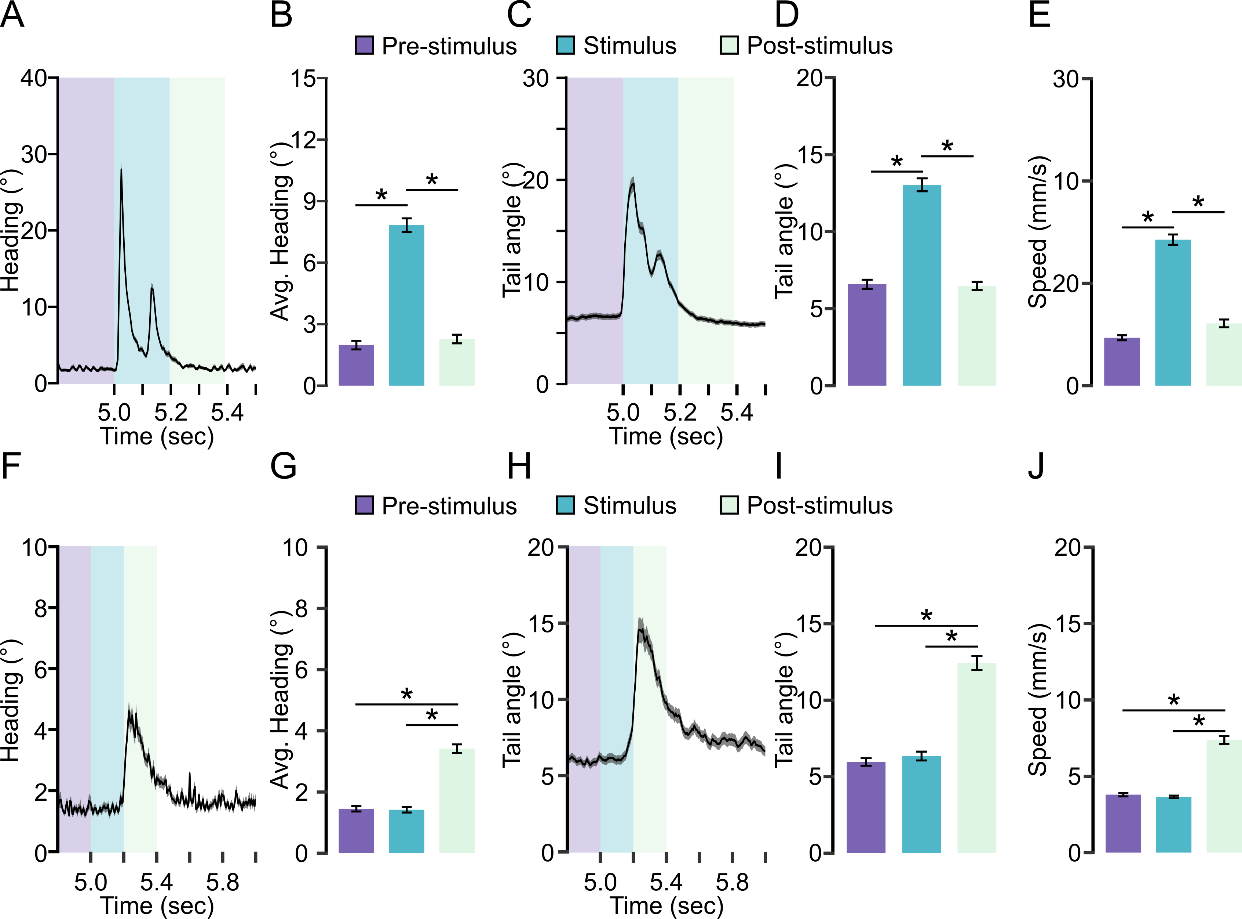

**Supplemental Figure 1 A-E.** Metrics plotted from individuals given a tap stimulus at 5.0 seconds in 96 well plate recordings for **A.** Average maximum absolute instantaneous heading change (N=643). **B.** Interval averaged maximum change in absolute heading direction **C.** Average maximum change in caudal tail angle **D.** interval averaged maximum change in caudal tail angle, or **E.** interval averaged maximum change in instantaneous speed. **F-J.** Same as in **A-E.** for individuals given a visual light off stimulus for 2 seconds (N=766). Color indicates interval (purple: 200 msec prior to onset of stimulus, blue: 200 msec immediately after onset of stimulus, green: 200 msec following the stimulation interval). * Indicates p < 0.05.

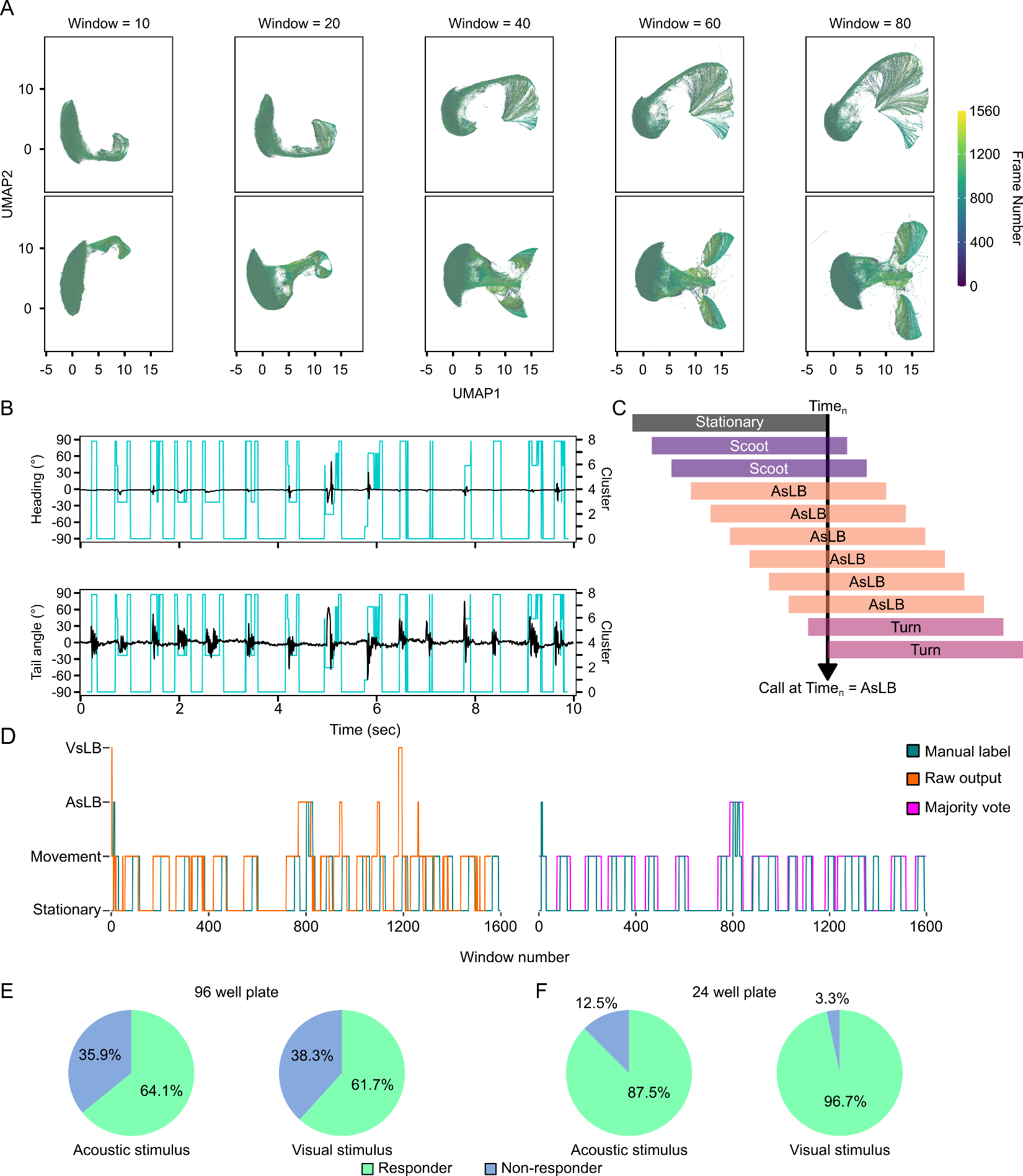

**Supplemental Figure 2 A.** UMAP representation of various window lengths. Color indicates the sequential order of window. **B.** Representative unsupervised k-means clusters (k=9, blue) overlaid on heading angle change (top) and instantaneous speed (bottom) for one fish. **C.** Schematic diagram of window aggregation method where at each timepoint all windows covering that timepoint are aggregated together and whichever window exhibits the majority call becomes the behavioral call for that timepoint. **D.** Comparison of raw output from model labeleing of a representative fish in a 96 well plate (orange) versus majority voting output (magents) compared to manual labeling of behaviors (teal). **E.** Responsive fish based on the presence of a call during the stimulus period across 6 acoustic startle acquisitions in 96 well plate recordings for acoustic and visual stimulation. **F.** Same as in **E.** for a VsLB call during the stimulus period for individuals recorded in the 24 well plate configuration.

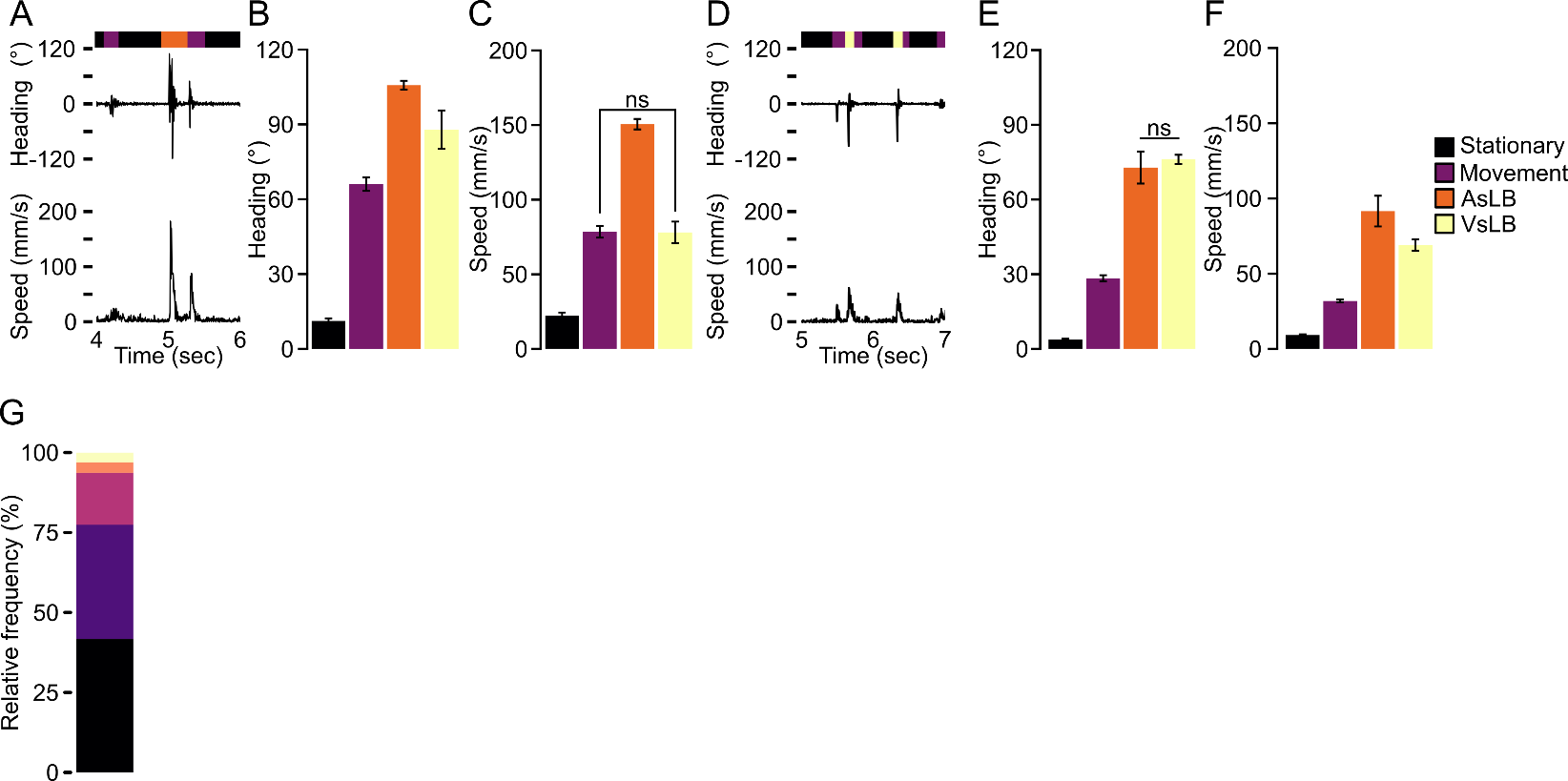

**Supplemental Figure 3 A.** Representative heaving angle change (top) and speed (bottom) aligned with behavioral calls (colored bar) for a 96 well plate individual during acoustic startle. Color indicates behavioral call type **B.** Average maximum heading angle change (left) or speed (right) per behavioral call type (Stationary: N=569, Movement: N=319, AsLB: N=412, VsLB: N=47) **C.** Same as in **B** for maximum instantaneous speed. **D-F.** Same as in **A-C** for visual startle (Stationary: N=384, Movement N=252, AsLB: N=30, VsLB: N=227). **G.** Relative frequency of call type among larvae including stationary calls. ns indicates p >0.05.

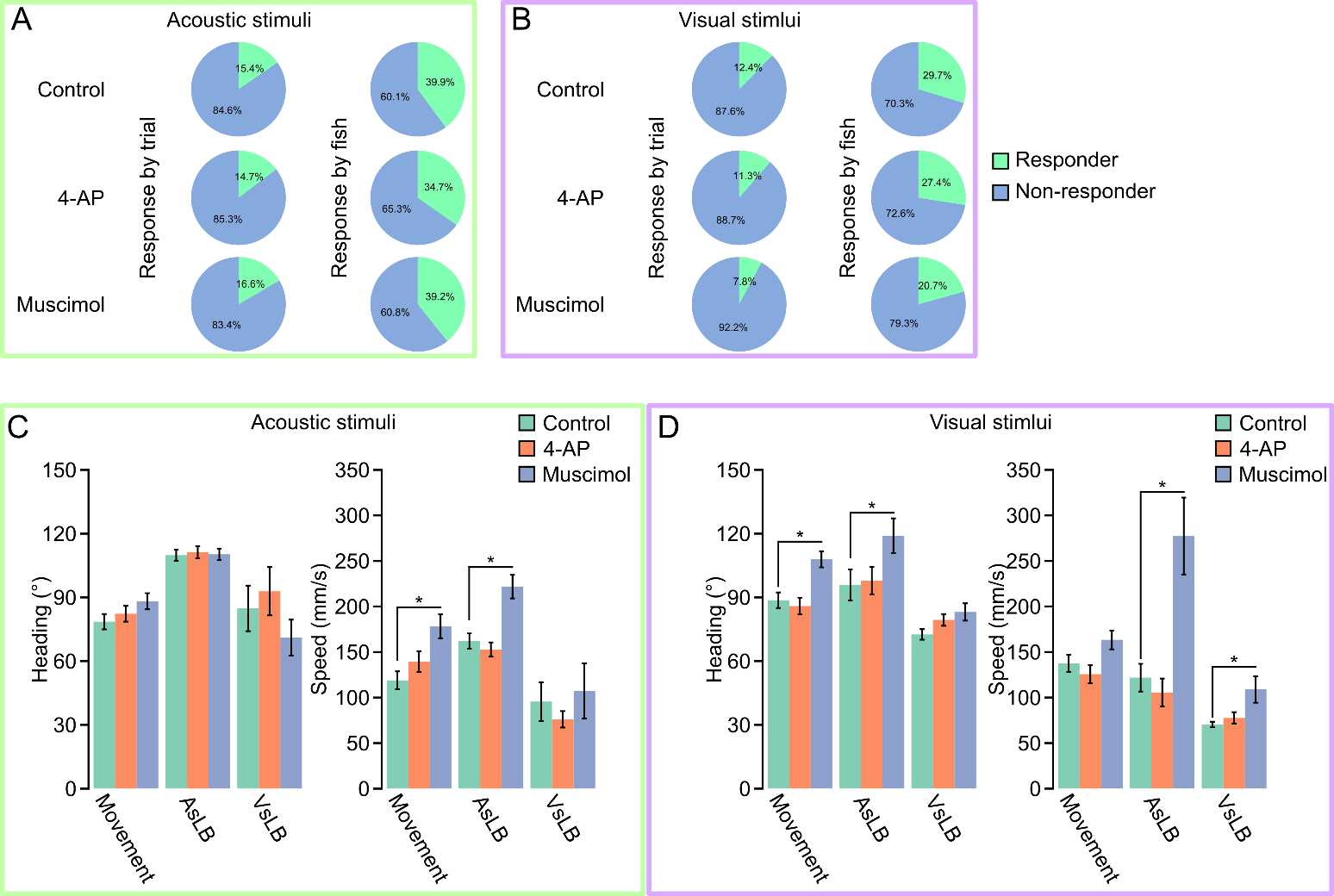

**Supplemental Figure 4** Proportion of responsive fish or responsive trials indicating one behavioral call for AsLB for **A.** acoustic startle (green outline) or **B.** VsLB for visual startle (purple outline) across 6 recordings for each drug treatment. **C.** Average maximum change in absolute heading direction or speed for each call type during acoustic stimulation (Control: Movement: N=298, AsLB: N=209, VsLB: N=32; 4-AP: Movement: N=246, AsLB: N=186, VsLB: N=18; Muscimol: Movement: N=286, AsLB: N=198, VsLB: N=30). **D.** Same as in **C.** for visual startle (Control: Movement: N=354, AsLB: N=53, VsLB: N=164; 4-AP: Movement: N=204, AsLB: N=50, VsLB: N=149; Muscimol: Movement: N=328, AsLB: N=46, VsLB: N=111).

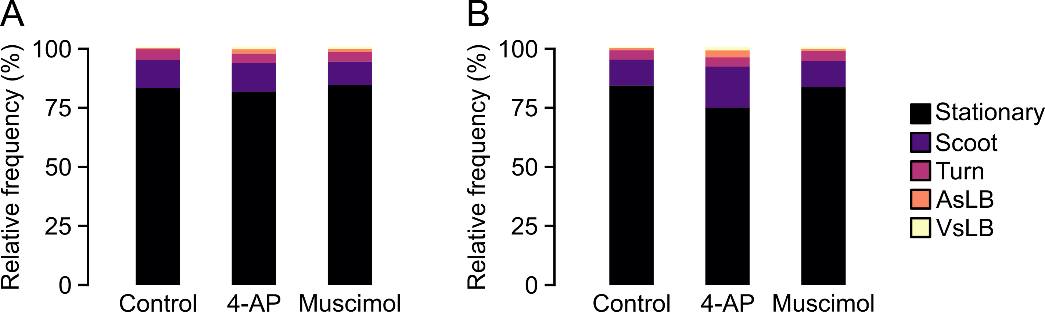

**Supplemental Figure 5 A.** Relative frequency of behavioral call type among drug treated larvae from 2-4 dpf including stationary calls. **B.** Relative frequency of behavioral call type among drug treated larvae tested in drug including stationary calls. Color indicates behavioral call type.
